# Narrow oviposition preference of an insect herbivore risks survival under conditions of severe drought

**DOI:** 10.1101/2020.03.24.005686

**Authors:** Ana L. Salgado, Michelle F. DiLeo, Marjo Saastamoinen

## Abstract

1. Understanding species’ habitat preferences are crucial to predict organisms’ responses to the current climate crisis. In many insects, maternal habitat selection for oviposition essentially determines offspring performance. Whether changes in climatic conditions may pose future mismatches in oviposition preference and offspring performance when mothers continue to prefer microhabitats that now threaten offspring survival is an open question.
2. To address this, we tested if oviposition preferences put offspring at risk in the Glanville fritillary butterfly (*Melitaea cinxia*) under drought stress. Mainly, we focus on identifying the microhabitat determinants for oviposition and the variation of conditions experienced by the sessile offspring, using field observations from 12 populations collected over 2015-2018. These data are combined with ten years of larval nest and precipitation data to understand within-population patterns of habitat selection. We tested whether the preferred microhabitats maximized the extended larval performance (i.e. overwinter survival).
3. We found that females preferentially oviposited in microhabitats with higher host plant abundance and higher proportion of host plants with signs of drought stress. In most years, larval nests had higher survival in these drought-stressed microhabitats. However, in an extremely dry year, only two nests survived over the summer.
4. Our results highlight that a failure to shift habitat preference under extreme climate conditions may have drastic consequences for the survival of natural populations under changing climatic conditions.

## INTRODUCTION

In the last century, ecosystems have experienced high rates of anthropogenic pressure including changes in climate, causing species to shift their ranges and/or significantly altering and even collapsing communities and ecosystems (Otto, 2018). Individuals require a specific habitat with the ideal abiotic and biotic conditions for development, reproduction and survival (Hanski, 2005; Kaweski, 2008). Examining how individuals select their habitat and how selection varies under different climatic conditions is essential for assessing species vulnerability under global change (Martin, 2001; Mayor, Schneider, Schaefer, & Mahoney, 2009). While many studies have highlighted the importance of habitat suitability for occupancy at the patch-scale, it is increasingly recognized that within-patch variation can play an important role in the persistence of populations (Hanski, I, 2005; Mortelliti, Amori, & Boitani, 2010). In particular, microhabitat variation can buffer populations from extreme conditions by providing refuges that allow individuals to remain under physiologically tolerable levels. However, this buffering effect is only possible if individuals are mobile and can track suitable conditions, or if a population exhibits variation in microhabitat use (Scheffers, Edwards, Diesmos, Williams, & Evans, 2014).

The factors that will determine if an individual accepts a habitat or not often depends on the environmental conditions, available resources as well as factors related to intra- and interspecific interactions (Morris, 2003; Schultz, Franco, & Crone, 2012). Many taxa have narrow microhabitat requirements, resulting in the use of only a small subset of potential available habitats. For example, in herbivorous insects, habitat use is tightly linked to local climatic conditions, quantity and quality of host plants, and the presence of associated species (e.g.: predators, parasitoids; Albanese, Vickery, & Sievert, 2008). For ectotherms that are sensitive to ambient temperature for thermoregulation, behavioural adjustment to microclimates (e.g.: moving short distances or ovipositing eggs in specific microclimates) is crucial. In temperate regions, warmer microclimates generally increase performance and survival (Caillon, Suppo, Casas, Arthur Woods, & Pincebourde, 2014; Derhé, Moss, Edwards, Carmenta, & Hassall, 2010; Scheffers et al., 2014). However, although average warming might benefit some temperate species (Caillon et al., 2014; Derhé et al., 2010; Roy & Thomas, 2003), increased extreme weather events can quickly push these warm microclimates beyond tolerable limits and consequently underlie some of the recently reported declines in insect abundances worldwide (e.g. Grubisic, van Grunsven, Kyba, Manfrin, & Hölker, 2018; Hallmann et al., 2017; Leather, 2018; Seibold et al., 2019).

In many insects, including butterflies, early larval stages are sessile; thus, the maternal oviposition choice determines the site where offspring will develop (Gripenberg, Mayhew, Parnell, & Roslin, 2010; Janz, 2003). Consequently, maternal oviposition site selection is of great importance for larval survival in specific microhabitats, considering that they will be restricted to the environmental conditions and the food quality of the host plant where they hatch (Martin, 2001; Schultz et al., 2012). Thus, the maintenance of maternal oviposition choice is linked to the selection pressure that offspring experience during development, and maternal fitness is linked to habitat elements that promote a better performance of her offspring (Albanese et al., 2008; Mayor et al., 2009; Rausher, 1979; Tjørnløv, Kissling, Barnagaud, Bøcher, & Høye, 2015). Many studies have found that maternal oviposition choice can enhance offspring fitness (preference-performance hypothesis; Gripenberg et al., 2010; Heisswolf, Obermaier, & Poethke, 2005; Wise & Weinberg, 2002). However, conflicts between maternal oviposition choice and larval performance may also arise (Janz, 2003; Mayhew, 2001). Firstly, what might be best for the offspring may not be the most suitable oviposition site for the female. Additionally, females divide their time between different tasks, which may create a conflict between the decision-making and the allocation of time and accuracy to oviposition and feeding, potentially leading to sub-optimal decisions on egg laying (Gripenberg et al., 2010; Janz, 2003). Finally, adults and larvae of many insects often feed on different host plant species or plant tissues (e.g.: nectar and plant tissue, respectively), or adults and larvae may have different nutritional requirements (Janz, 2005; Mayhew, 2001; Nestel et al., 2016).

Considering that it has increased its frequency and intensity in the last 50 years, drought is one climatic pattern that poses exceptional risk to individuals and populations in nature (Cook, Mankin, & Anchukaitis, 2018; Dai, 2011). For herbivorous insects in particular, drought can alter the quantity and quality of host plants where the offspring develop as water availability modifies host plant suitability (Albanese et al., 2008; John N. Thompson, 1988). We hypothesise that maternal habitat preference and offspring performance could be disrupted during extreme climatic events such as drought, i.e. when the mothers prefer to oviposit eggs in warmer and/or drier microhabitats. Such disruption may be especially likely at higher latitude where extremely hot and dry microhabitats may be favoured due to limited thermal conditions (Roy & Thomas, 2003). In previous studies, anthropogenic climate change has been linked to changes in the oviposition behaviour of the females, where the time and rate of oviposition and the range of suitable thermal locations has been altered (Davies, Wilson, Coles, & Thomas, 2006; Roy & Thomas, 2003). Most of the studies assessing impact of climate change on site selection so far have focused mainly on the direct effects (i.e., temperature), and thus neglecting the more indirect effects such as drought and the consequent changes in host plants.

We use the Glanville fritillary butterfly (*Melitaea cinxia*) and its metapopulation in the Åland islands, SW Finland, to assess within patch variation in habitat characteristics, such as host plant abundance and microclimatic conditions, and to understand microhabitat determinants for maternal oviposition choice. We capitalize on the systematic long-term monitoring data of the metapopulation (e.g. Hanski et al., 2017; Schulz, Vanhatalo, & Saastamoinen, 2019) and combine it with detailed field assessments to ask whether oviposition preferences vary across years and whether mothers generally choose locations that enhance their offspring’s overwinter survival. As Finland is the northern range limit of these butterfly species, with relatively short time window for larval development, and based on our recent findings that larval performance may be increased by feeding on drought-exposed host plants (Rosa, Minard, Lindholm, & Saastamoinen, 2019; Salgado & Saastamoinen, 2019), we hypothesize that drier microhabitats are generally favoured by the females but may be disadvantageous for the larvae during particularly warm and dry years.

## MATERIALS AND METHODS

### Study system

The Glanville fritillary inhabits fragmented landscapes in the Åland Islands (Finland), where it persists as a classical metapopulation in a ∼4500 meadow (habitat patch) network. Females frequently disperse among nearby meadows in search of nectar and oviposition sites (DiLeo, Husby, & Saastamoinen, 2018; Niitepõld, Mattila, Harrison, & Hanski, 2011). In early June, mated females lay 100-200 eggs in batches on the host plants *Plantago lanceolata* or *Veronica spicata*. The larvae hatch in late June and early July and feed gregariously on the host plant where they were deposited by their mother. The small larvae are sessile and can move only short distances (i.e. < 1m) after defoliating their host plant (Kuussaari, Van Nouhuys, Hellmann, & Singer, 2004). The larvae overwinter as a group, mostly as fifth instar within a dense silk web. In the next spring they break diapause and start feeding on the new host plant growth to complete two more larval instars before pupation (Saastamoinen, Hirai, & van Nouhuys, 2013). The weather conditions experienced throughout the lifecycle impact the population dynamics of the butterfly (Kahilainen, van Nouhuys, Schulz, & Saastamoinen, 2018).

The habitat patches generally are very small (mean = 1500 m^2^) but characterised by high host plant density that can support several larval groups. Even though large patches may have a lower density of host plants, they often have high number of nectar plants to sustain higher number of adults (Nieminen, Siljander, & Hanski, 2004). The severity of host plant drought prevalence assessed during the fall survey of the butterfly is known to vary across the habitat patches (Hanski et al., 2017). This variation is likely caused by several reasons such as variability in local weather conditions and types of habitat patches (e.g. pasture vs. outcrop meadow) as well as by more human induced changes such as grazing or presence of roads. Additional variation at smaller spatial scales within patches occur due to soil type, rocks or slopes, creating specific microclimatic conditions (Kuussaari et al., 2004). In general, summer drought has increased in frequency during the last decades in the archipelago (Tack, Mononen, & Hanski, 2015). Previous studies have shown that food scarcity and high levels of desiccated host plants at the end of the summer may lead to reduced body mass and even result in starvation of the larvae, which may consequently reduce overwinter survival and cause local extinctions (Kuussaari et al., 2004; Nieminen et al., 2004). Long-term monitoring of the metapopulation has led to a good understanding of the determinants of overall patch quality on the occupancy and abundance, however, little is known about within-patch variation (but see Schulz et al., 2019).

### Field observations and data collection

We studied 12 meadows with *P. lanceolata* as the only host plant. These habitat patches were located across the main island and were selected based on consistent presence of larval nests in 2012-2014 to minimize the risk of local extinction in the following years. For more information about the population selection please see Salgado et al. (in prep). Each of the selected populations was divided into a grid composed of 20×20 m cells that fell within the patch boundaries (see figure S1 in Supporting Information). For each cell we determined host plant abundance (proportion of the area) and the proportion of the host plants that showed signs of drought stress (i.e. wilting; details in Appendix S2). During 2015-2017, these measurements were determined from the end of June until the end of August four times per patch with 15 days interval between the subsequent surveys. An additional year was added in 2018 due to an extreme summer drought. In this year we collected the host plant abundance and site drought stress information only once (July 13^th^-18^th^). In 2016 and 2017, we added information of the vegetation structure of the cells. These data included: vegetation height (m), canopy coverage (%) and abundance of nectar plants (%). We included the topography of the cells: soil depth (cm), terrain slope (degrees) and terrain aspect (degrees). Using the terrain aspect, we extracted values for the transformed field aspect (tasp) and the slope corrected transformed aspect (tasl; details in Appendix S2; Lookingbill & Urban, 2004). Although we were unable to directly follow temperature in each cell, local thermal conditions should, in part, be captured by measures of vegetation structure and topography.

Presence and number of nests per grid cell were taken from long-term autumn surveys that have occurred annually over the last 28 years (Ojanen, Nieminen, Meyke, Pöyry, & Hanski, 2013). Every autumn, a group of field assistants record the status of the metapopulation, and from 2009-2018 GPS locations of all larval nests were recorded. We used QGIS version 2.12.0 (http://www.qgis.org/) to count the number of nests per cell in each of the populations from 2009-2018. To confirm that locations of nests in the autumn surveys reflect the initial oviposition locations, we compared occupancy models using the autumn nest locations and locations from a smaller survey in 2015-2017 that followed nests at early stages (eggs-fourth instar) and recording locations at the end of summer, and found that they did not differ (details in Appendix S2). To be able to include data from 2018 in our models, nest locations observed during the summer were used instead of the autumn data. No permissions were required for performing the present study.

### Data analyses

All statistical analyses were conducted in R 3.5.1 (R Development Core Team, 2018). For all analyses, we only included cells with at least 1% host plant coverage. To reduce the number of covariates in our models, we conducted a principal component analysis (PCA) on variables describing cell topography and vegetation. Variables included in the PCA were vegetation height, canopy coverage, soil depth, tasp, and tasl (described above and in Appendix S2). Variables were centred and scaled prior to analysis, and principal components with eigenvalues greater than one were retained for downstream analysis.

### Variation in host plant abundance and drought stress

Our first aim was to quantify microhabitat variability to understand the range of local conditions. We used variance partitioning on generalized mixed models to separate sources of variation in host plant abundance and drought stress measured across the different time points (i.e. four repeated measures per year) into the nested sampling scales of year > population > cell. Because host plant abundance and drought stress within each site were measured as proportions and data were unbalanced, we estimated variance components from beta mixed models implemented in the glmmTMB library (details in Appendix S2; Brooks et al., 2017). We further calculated pairwise correlation coefficients between measurements of site’s host plant abundance or proportion of host plants with signs of drought stress taken across different time points and years.

### Testing for oviposition site preference

We used generalized linear mixed models with binomial error distribution to test if females preferential oviposit in microhabitats of high or low host plant abundance and in microhabitats experiencing high or low levels of drought or other environmental variables. The presence (1) or absence (0) of a nest in each cell per year (2009-2018) was entered as the response variable. For each cell, we summarized the host plant abundance and the proportion of the host plants with signs of drought stress by taking the mean of field measurements across the four years it was measured, 2015-2018. The mean proportion of host plants with signs of drought stress was calculated by averaging values measured during the driest summer period per year (time point four in 2015, and time point three in 2016 and 2017, only one time point was measured in 2018), as this explained the most variation in the data compared to values from single time points or the mean or maximum value. We thus take the mean proportion of host plants with signs of drought stress to reflect how prone a site is to dry out during the summer. We acknowledge that these averaged values may not necessarily reflect the drought conditions experienced during the oviposition period. However, as plant phenotypes change within a matter of days to weeks in response to drought stress (Salgado A.L. personal observation), we believe that the averaged values are more representative of the microsite’s stress to drought than measures from single snapshots in time. We further include an interaction with weather conditions during the oviposition period to better reflect actual drought experienced (details below). Finally, as a further test of our decision to average drought stress values over the four-years, we compared our results to a model that included drought stress and host plant coverage per year for the years the data were available (2015-2018). After removing cells with missing values, we had a final sample size of 1403 presence/absence and abundance observations for the years 2009-2018.

The proportion of abundance of nectar plants and the first three principal components of our PCA describing site topography and vegetation were also included as covariates in the model. To test if weather during early summer (i.e. immediately before and during the oviposition period) influences oviposition decisions, we additionally included two-way interactions between May and June precipitation and host plant variables. Monthly precipitation amounts for May and June were extracted for each cell in each year from 10×10 km gridded data provided by the Finnish Meteorological Institute (Aalto, Pirinen, & Jylhä, 2016). Fixed covariates were centred, scaled and were tested for collinearity prior to inclusion. To account for non-independence of data collected from nearby cells and repeated measures from successive years, we included a spatiotemporal random effect using Integrated Nested Laplace Approximation with Stochastic Partial Differential Equations (INLA-SPDE; (Lindgren, Rue, & Lindström, 2011), implemented in the R INLA library (Rue, Martino, & Chopin, 2009). This method is increasingly being used in ecological studies as an efficient way to model species occurrences and population dynamics while accounting for spatial and temporal dependencies in the data (e.g.: Myer, Campbell, & Johnston, 2017; Schulz et al., 2019; Ward et al., 2015). In brief, spatial dependency of observations are accounted for using a latent Gaussian random field, which we constructed using a two-dimensional irregular grid (mesh) based on the geographic coordinates of cell centroids. Exploratory analysis indicated that spatial autocorrelation was present within but not between patches, and we thus constructed meshes within patches only to speed up computation time (see Figure S3). Temporal dependencies in the data were accounted for by including a residual autoregressive correlation of order one (AR1). We further wanted to understand the factors that predict the number of nests found in each cell. Because our data included an excess of zeros (i.e. nest absences), we used a hurdle model to jointly predict zero-inflation (i.e. nest presence/absence) and nest counts. Unlike zero-inflated Poisson models, hurdle models assume that zeros (i.e. nest absences) are real and are driven by the same processes that drive non-zero observations, which we believe to be true for our system. We implemented the hurdle as a two-component model that mixes binomial and Poisson distributions, with the first component predicting the binary outcome of nest presence or absence, and the second component predicting the positive counts of nests. We included the same covariates (see above for binomial model) in both components of the model, and additionally included the log of cell area as an offset in the Poisson component as some cells slightly deviated from the 20×20 m size.

### Effects on nest survival

For each year of the period 2009-2017, larval nests found in the autumn were visited again the following spring to quantify overwinter survival. We used these data to test if female oviposition preferences affected overwintering survival. We used a binomial model with the number of surviving nests per cell as the response variable (i.e., number of successes), weighted by the total number of nests found in the previous autumn (i.e., number of trials). Only cells that had nests in the autumn were included in the analyses, giving a final sample size of 271 observations. To accommodate this smaller sample size and avoid overfitting, we included only those covariates that were found to be important in our analyses of oviposition preferences and did not include interaction terms. Host plant abundance, proportion of host plants with signs of drought stress, PC1, and PC2 were included as fixed covariates, and we included a spatiotemporal random effect to account for the non-independence of the data.

## RESULTS

### Principal component analysis

The principal component analysis reduced the vegetation and topographic variables into three components with eigenvalues greater than one, explaining 66% of the total variation (see Table S4). The first principal component explained 28% of the total variation, with strong negative loadings of tasp and tasl (i.e. low values of PC1 reflect steeper, more south-western facing slopes; see Table S4). The second principal component explained 20% of the variation, with strong positive loadings of soil depth and canopy coverage. The third principal component explained 18% of the variation with strong positive loadings of vegetation height and negative loading of slope.

### Variation in host plant abundance and drought stress

Variance partitioning of host plant abundance indicated that most variation in the data was explained by cell (0.33) followed by population (0.22), with little variation in abundance among years (0.01; fig. 1; see Figure S5). In contrast, the component for cell for proportion of host plants with signs of drought stress was near zero, indicating that the measurements were not repeatable across time points. The component for patch was also low (0.05), indicating patches did not differ in mean drought stress. The majority of the variation (0.64) in drought stress could be explained by differences among years, with 2015 being a very wet year with low drought stress values and 2018 being an extremely dry year with high drought stress values (fig. 1). Although 2018 was a year of extreme drought, we still found substantial variation in drought stress among microsites within patches (see Figure S6).

**Figure 1.**
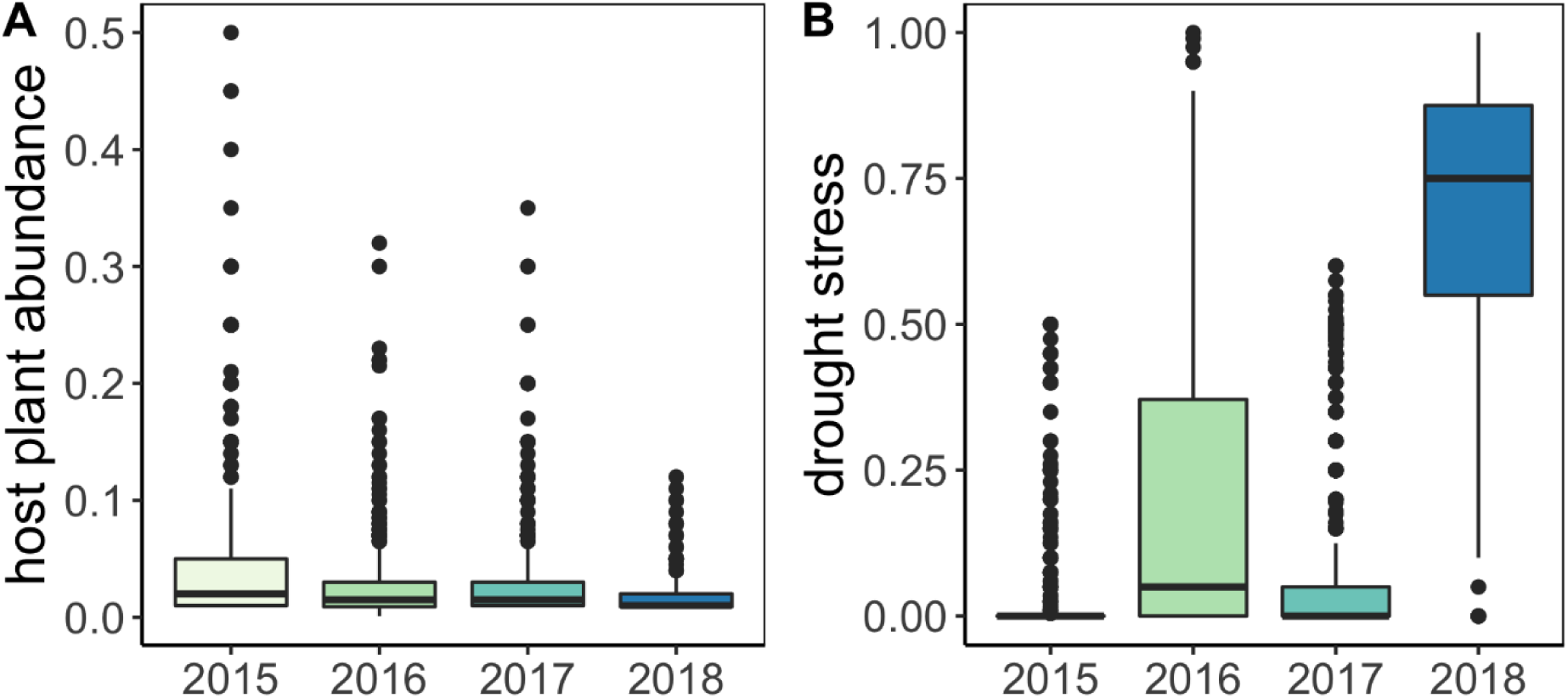
Boxplots showing variation in host plant abundance (A) and drought stress (B) in cells for 2015-2018. See S for variation among populations.

Measurements of host plant abundance showed consistently strong positive pairwise Pearson correlations among time points and years, with correlations between yearly means ranging from 0.6-0.8 (see Figure S7). Pairwise Pearson correlations between measurements assessing the proportion of host plants with signs of drought stress taken at different time points and years showed mostly positive but weaker correlations (yearly mean pairwise Pearson correlations from 0.2-0.6), with the exception of measurements taken in 2015 and 2017, which were negatively correlated with one another or showed no correlation (see Figure S8).

### Testing for oviposition site preference

All covariates were uncorrelated (see Figure S9) and had variance inflation factors below two, indicating that our variables capture different aspects of the microhabitat. The proportion of host plant abundance and the proportion of host plants with signs of drought stress were positively correlated with nest presence, while PC1 and PC2 were negatively correlated with nest presence (table 2, fig. 2). This indicates that nests are more likely to be found in sites of high host plant abundance and areas that are more prone to drought stress, on south-west facing slopes (PC1) and in shallow soils and open canopies (PC2). The model additionally showed positive effects of June precipitation on nest presence, and an interaction between drought stress and May precipitation. This suggests a higher probability of occupancy in wet years or wet patches, but also that nests were more likely to be found in drought-prone microsites in drier years (fig. 3). The credibility intervals of abundance of nectar plants and PC3 overlapped with zero, indicating that these variables were not good predictors of nest presence/absence. In the hurdle model, zero-inflation probability was explained by the same covariates as the binomial model (see Table S10). Host plant abundance, drought stress, June precipitation and PC3 were positively related to truncated Poisson nest counts, while PC2 was negatively related to nest counts (see Table S11). This indicates that a higher abundance of nests tends to be found in sites of high host plant abundance that are more prone to drought stress, and in shallow soils with open canopies (PC2) and flatter sites with higher vegetation (PC3).

**Table 2.** Posterior mean estimates, standard deviations (sd) and 95% credibility intervals from spatiotemporal binomial INLA models on nest presence using ten years of nest presence/absence observations. Parameters values in bold are significant.

| Parameter | mean | sd | 95% credibility |  |  |
| --- | --- | --- | --- | --- | --- |
| Intercept | **-2.1** | **0.26** | **-2.6** | , | **-1.6** |
| Host plant abundance | **0.75** | **0.13** | **0.51** | , | **1.00** |
| Drought stress | **0.72** | **0.16** | **0.41** | , | **1.04** |
| Abundance of nectar plants | 0.09 | 0.12 | -0.14 | , | 0.33 |
| PC1 | **-0.33** | **0.11** | **-0.56** | , | **-0.11** |
| PC2 | **-0.44** | **0.15** | **-0.73** | , | **-0.15** |
| PC3 | -0.01 | 0.14 | -0.29 | , | 0.27 |
| May precipitation | -0.08 | 0.10 | -0.27 | , | 0.11 |
| June precipitation | **0.31** | **0.10** | **0.11** | , | **0.51** |
| Host plant abundance x May precip | -0.08 | 0.08 | -0.23 | , | 0.07 |
| Drought stress x May precip | **-0.23** | **0.10** | **-0.43** | , | **-0.04** |
| Host plant abundance x June precip | 0.00 | 0.09 | -0.17 | , | 0.16 |
| Drought stress x June precip | 0.15 | 0.11 | -0.06 | , | 0.36 |
| Random effects |  |  |  |  |  |
| Temporal correlation | 0.93 | 0.03 | 0.85 |  | 0.98 |
| Variance of spatial random effect | 1.5 |  |  |  |  |

**Figure 2.**
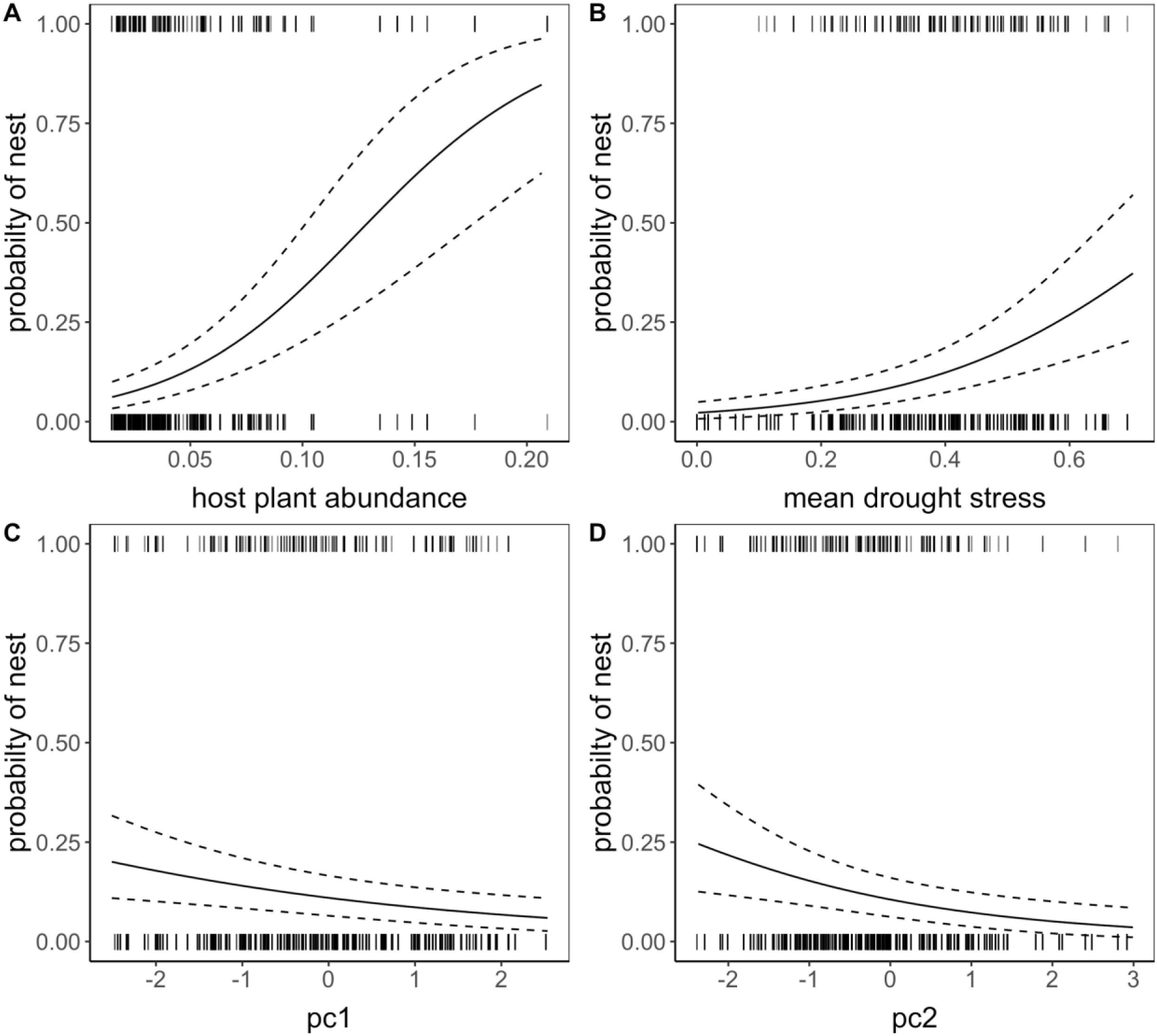
Relationships of host plant abundance (A), the mean proportion of host plants with signs of drought stress (i.e. drought stress; B), PC1 (C) and PC2 (D) with the probability of nest presence for 2009-2018. Vertical ticks at zero and one show nest absence and presence respectively. Solid black lines are prediction of fixed effects from the spatiotemporal binomial model and dashed lines show 95% credibility intervals.

**Figure 3.**
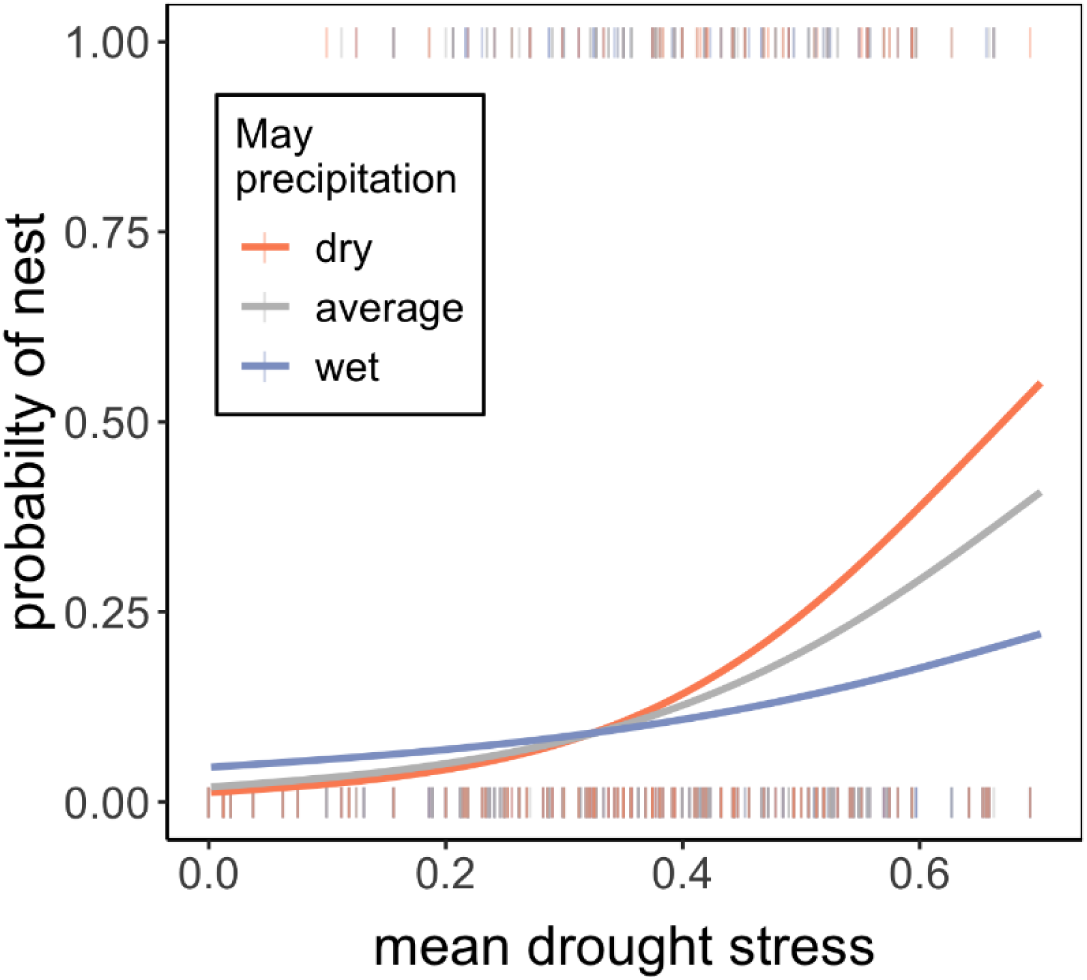
Relationship between mean proportion of host plants with signs of drought stress, May precipitation, and the probability of nest presence for 2009-2018. Solid line are predictions of the fixed effects from the model including an interaction between drought stress and May precipitation. May precipitation was divided in: dry years, which includes any precipitation values below one standard deviation of the mean; average years, which includes values within one standard deviation of the mean, and wet years, which includes values above one standard deviation from the mean. Vertical bars at zero and one are observed nest absences and presences, respectively.

Nest observations were highly correlated in time (temporal autocorrelation was 0.93 and 0.76 for binomial and hurdle model respectively; Table 2; see Table S11), suggesting that females tend to put nests in the same microsites year after year. Nest observations were significantly spatially autocorrelated (i.e. clustered) up to a range of 67 m in the binomial model, and 50 m in the hurdle model.

The same factors (host plant abundance, drought stress, PC1, and PC2) were identified as having significant effects on nest presence in the binomial model that included per-year measures of host plant coverage and drought stress using the four-years of data for which these measures were available (see Table S11). This suggests that averaging of drought stress and host plant abundance values in the long-term model had little influence on the results. We did, however, find some differences in estimates of nest abundance; namely, the four-year model found no significant effect of drought stress and PC2 on nest abundance and found a significant negative effect of PC1 (see Table S11). Models using summer nest locations, autumn nest locations, and a mix of autumn (2015-2017) and summer nests locations (2018) provided similar results (see Table S11), suggesting that our use of autumn nest locations from the long-term survey data can be taken to reflect nest locations during the oviposition period.

### Effects on nest survival

Overwintering survival was high, with 72% of nests surviving winter over 2009-2017, and yearly survival rates ranging from 57-85%. The spatiotemporal binomial model indicated that the probability of overwinter survival was positively related to the proportion of host plants with signs of drought stress in the cell (table 3; fig. 4). In comparison, all other covariates had credibility intervals overlapping zero, indicating that they were not good predictors of overwinter survival.

**Table 3.** Posterior mean estimates, standard deviations (sd) and 95% credibility intervals from binomial INLA model on overwinter survival using nest presence/absence data (2009-2017). Host plant abundance and the proportion of host plants with signs of drought stress represent mean values across four years of field measurement per cell (2015-2018). Parameter values in bold are significant.

| Parameter | mean | sd | 95% credibility interval |  |
| --- | --- | --- | --- | --- |
| Intercept | **1.4** | **0.23** | **0.95** | , **1.9** |
| Mean host plant abundance | -0.15 | 0.15 | -0.44 | , 0.15 |
| Mean drought stress | **0.31** | **0.15** | **0.01** | , **0.62** |
| PC1 | -0.06 | 0.15 | -0.35 | , 0.23 |
| PC2 | -0.06 | 0.16 | -0.39 | , 0.25 |

**Figure 4.**
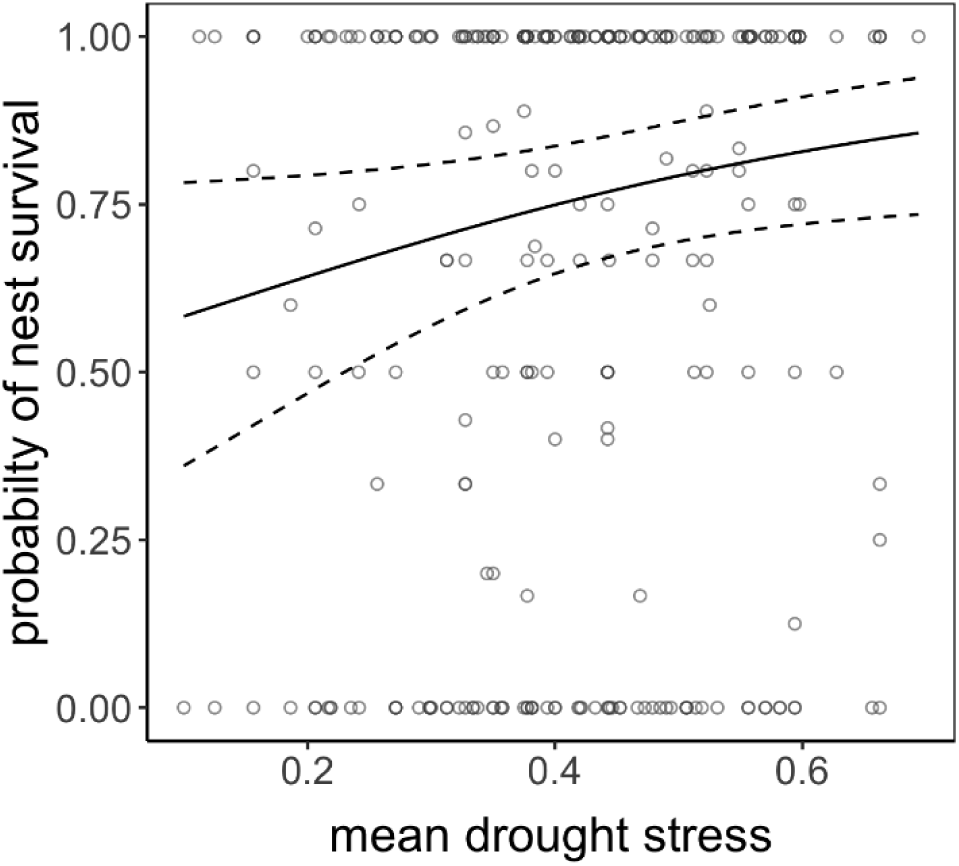
Relationship between the mean proportion of host plants with signs of drought stress and overwinter nest survival for 2009-2017. Points are observed proportions of surviving nests per cell. The solid black line is the prediction from a spatiotemporal binomial INLA model. Dotted lines show the 95% credibility interval.

## DISCUSSION

In this study, we identified the determinants of oviposition site choice, and assessed its temporal and spatial variation in a temperate butterfly species. We were specifically addressing the correlation between maternal habitat preference and offspring performance and whether changing climatic conditions may disrupt this correlation. Our results show that female preference of oviposition for microhabitats with higher host plant abundance and higher proportion of host plants with signs of drought stress increases offspring survival in normal years. However, the failure of females to shift their preferences during the summer of intense drought resulted in a dramatic decrease in offspring survival.

### Variation in host plant abundance and drought stress

Variance partitioning showed that most of the variation in host plant abundance was within populations, observed as high variation among cells. Meanwhile, considerable variation in the proportion of host plants showing signs of drought stress occurred between years (fig. 1). In contrast, the variance component of drought stress for cell was near zero, however, this does not indicate a lack of variation. Rather, we found that drought stress varied so much between the four measured time points per year that there was no difference in cell means. This makes sense considering that drought stress values tended to be low or zero during wetter time periods and high during drier periods. Taking values from only the driest time period per year, we observed considerable variation among cells within populations (see Figure S6). Together our results suggest that the habitats of the Glanville fritillary in Åland vary in space and time, with high fluctuations in the quality of host plant over space and time, and high variation in the quantity of the host plants in space. Furthermore, plant populations can vary in age and size (not assessed), which can further impact the interactions with herbivores and their hosts (Thompson, 1988). At fine scales, habitat structure and topography can be essential in determining both resource availability and the microclimatic conditions (Wilson et al., 2015).

### Testing for oviposition site preference and effects on larval survival

We further established the factors that determine habitat selection for oviposition. Our models showed that host plant abundance, drought stress, PC1 and PC2 predicted the presence of the nests within populations, which can be linked to maternal oviposition choice. The oviposition preference within the habitat increased with a higher proportion of host plant abundance and a higher proportion of host plants with signs of drought stress. Meanwhile, it increased with a higher transformed field aspect (tasp) and slope corrected transformed aspect (tasl; reflected in PC1), and decreased with canopy coverage and soil depth (PC2), suggesting that females prefer to oviposit on south-facing slopes and in open microhabitats with shallow soils, respectively. While host plant abundance and PC1 predicted the number of nests in the model including four-years of data, host plant abundance, drought stress, and abiotic microhabitat characteristics (PC2 and PC3) predicted the number of nests found in microhabitats using the long-term data. This discrepancy indicates that certain factors were only important in some years, or that their effects on the number of nests are small and only detectable with more data. Nevertheless, our results indicate that different factors are important for determining nest presence versus nest abundance.

Our results are in hand with previous observations that ovipositing herbivorous insects aggregate in areas with high abundance of host plants and can further select sites based on the quality of the microhabitat (Janz, 2003). In the UK, the distribution of larval groups of the Glanville fritillary are similarly restricted to warmer areas, as the mothers prefer host plants that are warmer than the ambient temperature (Curtis & Isaac, 2015). This choice of warmer microhabitat can increase the performance of offspring by helping with thermoregulation and increasing the metabolism, resulting in faster developmental times (Curtis & Isaac, 2015). We found that, in all years except 2018, nests in microhabitats that were more prone to drought stress were more likely to survive overwinter. Interestingly, of all the factors tested, only the drought stress affected overwinter survival, and host plant abundance was not an important predictor despite the strong preference for ovipositing in areas of high host plant abundance. The fact that host plant abundance was not important could indicate that there is no limitation of food resource within these populations, and that even in microhabitats with low host plant abundance there is enough host to feed on. Considering the lifecycle of the butterfly, the larval instars during the summer are still relatively small, and even though they live gregariously they rarely consume large number of plants at this stage. The maternal choice of high host plant abundance sites may also reflect the offspring needs in the following spring. At this stage the postdiapause larvae consume more host plants and often run out of food (Saastamoinen M. personal observation).

Our finding that female choice and larval survival was positively linked to how prone a site is to drought stress in the field, may simply result from these microhabitats being also warmer and thus inducing faster larval development (Roy & Thomas, 2003). However, several lines of evidence support an important role for drought stress over local thermal conditions. First, although we did not measure temperature of microsites directly, we did measure several aspects of topography and vegetation that should reflect local temperatures (e.g. low values of PC1 and PC2 reflected southwestern facing slopes with open canopies). These measures were uncorrelated with our measure of drought stress, suggesting that they fundamentally capture different aspects of the microhabitat or microclimate. Second, while we did find that females prefer to oviposit in microsites associated with warmer thermal conditions (i.e. nest presence showed significant negative relationships with PC1 and PC2) in addition to sites with higher host plant coverage and drought stress, neither PC1 nor PC2 were found to be related to overwintering survival. Finally, we have some experimental evidence that suggests that feeding on drought exposed host plants does, at least in some families, directly increase larval performance (pre-diapause larvae: Rosa et al. 2019 & post-diapause larvae: Salgado & Saastamoinen, 2019) that seems to even translate to increased adult performance (Salgado & Saastamoinen, 2019). Even though the impact of drought on host plant quality was not assessed here, previous studies on other systems have shown that plants under drought stress often accumulate nutrients, such as carbon and nitrogen, that can enhance the performance of insect herbivores (Gutbrodt, Mody, & Dorn, 2011; Mattson & Haack, 1987). We note however, that as previous laboratory studies (e.g. Ahola et al., 2015; Kallioniemi & Hanski, 2011; Kvist et al., 2013) in *M. cinxia* have shown temperature to play a central role in larval development, as expected for any ectotherm, it is also possible that we failed to find this relationship in the field because our topographic and vegetation variables capture only a part of the local thermal conditions. Therefore, further experiments are required to validate the specific aspects of host plant quality and abiotic conditions that serve as cues for oviposition and that affect offspring survival.

We found that weather conditions played an important role in predicting the presence and number of nests. We found a positive relationship between June precipitation and nest presence and count, suggesting that while females prefer dry microhabitats, they also tend to lay more clutches in wetter patches or years. This could be linked to the physiological needs of the larvae. Kahilainen et al. (2018) showed that growth rates of populations were strongly positively correlated with spring precipitation, indicating that moist conditions are important for larval development. We note that in most years, patches that received more rain in June were also warmer on average (see Figure S12), which might suggest that a more complicated combination of temperature and moisture determines ideal conditions for oviposition, egg hatching rates or larval survival at early stages of development. While we found no evidence of a direct effect of May precipitation on nest presence or count, May precipitation interacted with drought stress to influence oviposition choices. Specifically, the positive relationship between nest presence and whether the host plants within the site were prone to drought was strongest in the driest years and weakest in wet years. A potential explanation for this is that host plants do not experience drought stress under wet conditions leading to low spatial variability within populations, and thus females have no opportunity to be choosy. May precipitation appears to be important because it impacts food abundance and quality of host plants in the next months, which are crucial for the development of the prediapause larvae. Another possibility is that females can use May precipitation as a predictor of host plant quality in the future months.

Crucially, our results showed that females did not shift their preferences in times of extreme drought (i.e. 2018). This is surprising as by the time mothers were ovipositing, the host plants were extremely dry due to unusual weather conditions in May 2018 (i.e. high thermal conditions combined with extremely low precipitation levels; van Bergen et al., in review). This may suggest that the females lack the ability to shift their microhabitat preferences. Previous studies of *Hesperia comma*, on the contrary, have shown that females can adjust their preferences according to the climatic conditions experienced, as warmer host plants are selected for oviposition at low temperatures, and cooler host plants at high temperatures (Davies et al., 2006). These shifts are important for tracking optimal conditions for offspring and buffering the effects of climate change (Scheffers et al., 2014). For example, in warblers, the location of nests has shifted as a result of altered long-term precipitation patterns (Martin, 2001).

The failure of females to shift their preferences during the summer drought of 2018 had drastic results, with only two larval nests surviving to autumn (instead of 87, 88, and 54 in the previous years). The two surviving nests were found in a cell that had a mean drought stress value within the lowest quartile of all microhabitats used by females that year (see Figure S13), highlighting that microhabitat can play an important role in buffering populations from extreme events, but only when accompanied by enough variation in oviposition preferences (Derhé et al., 2010; Scheffers et al., 2014). If flash drought, which appears with no warning and intensifies rapidly within a season (Cook et al., 2018), starts to become more and more common in the next years, the entire metapopulation could be at risk. If the maternal oviposition preferences are heritable then it is possible that the extreme drought could select for females that prefer to oviposit in slightly moister areas, which will contribute for a greater development and survival of the offspring (Thompson & Pellmyr, 1991). The observed dynamics are evidence of the importance of the interaction between abiotic and biotic factors on habitat selection and the implications for the species and their ecological consequences under novel environmental conditions (Martin, 2001).

## CONCLUSIONS

This study shows that females of the Glanville fritillary have strong oviposition preferences linked to microhabitats with high host plant abundance and proneness to harbour drought stressed host plants. In most years, these preferences appear to be adaptive as larval nests in drought prone microhabitats were more likely to survive overwinter. However, with only two nests surviving in a year of extreme drought, our results suggest that this preference-performance can be disrupted by extreme climatic events. Sudden and unpredictable alterations in environmental conditions (e.g. temperature and precipitation) that consequently impact strategies evolved to fine-tune maximization of individual’s performance can thus have devastating consequences. With drought becoming more frequent and severe, number of species could be at risk because of insufficient plastic responses (Caillon et al., 2014; Cook et al., 2018; Roy & Thomas, 2003) allowing shifts in strategies that have become maladapted due climate change. Such lack of variation in site preference may even play a role in the recently reported insect declines worldwide.

## ACKNOWLEGEMENTS

We thank to our field assistants Elina Laurén, Heini Karvinen, Alma Oksanen, Juha-Matti Pitkänen, Susanna Rokkanen and Paula Salonen for their help in data collection during the summers, also to all the field assistants that worked on the autumn surveys (2009-2018). Many thanks to Aapo Kahilainen for helping on the patch selection and advice on analysis, to Torsti Schultz for providing part of the topography data, and to Erik van Bergen for comments, as well for the contribution of two anonymous reviewers. The research was funded by European Research Council (independent starting grant no. 637412 ‘META-STRESS’ to MS). We affirm that we have no conflict of interest.

## AUTHORS CONTRIBUTIONS

MS and ALS conceived the present idea and planned fieldwork; ALS carried data collection; MFD performed data analyses; ALS and MFD wrote the first draft of the manuscript; All authors provided critical feedback and helped to shape the research, analyses, manuscript gave final approval for publication.

## DATA AVAILABILITY STATEMENT

Data in Dryad Digital Repository upon acceptance.

