## Supplemental Information for "Narrow oviposition preference of an insect herbivore risks survival under conditions of severe drought"

SUPPORTING INFORMATION

Figure S1. The main islands of Åland with the 12 populations selected for data collection during summers 2015-2018. Population 1621 (right down panel) shows the 20x20 grid (purple lines) superimposed to divide the habitat in cells and to collect detailed vegetation structure of each cell within the patch borders (red line).

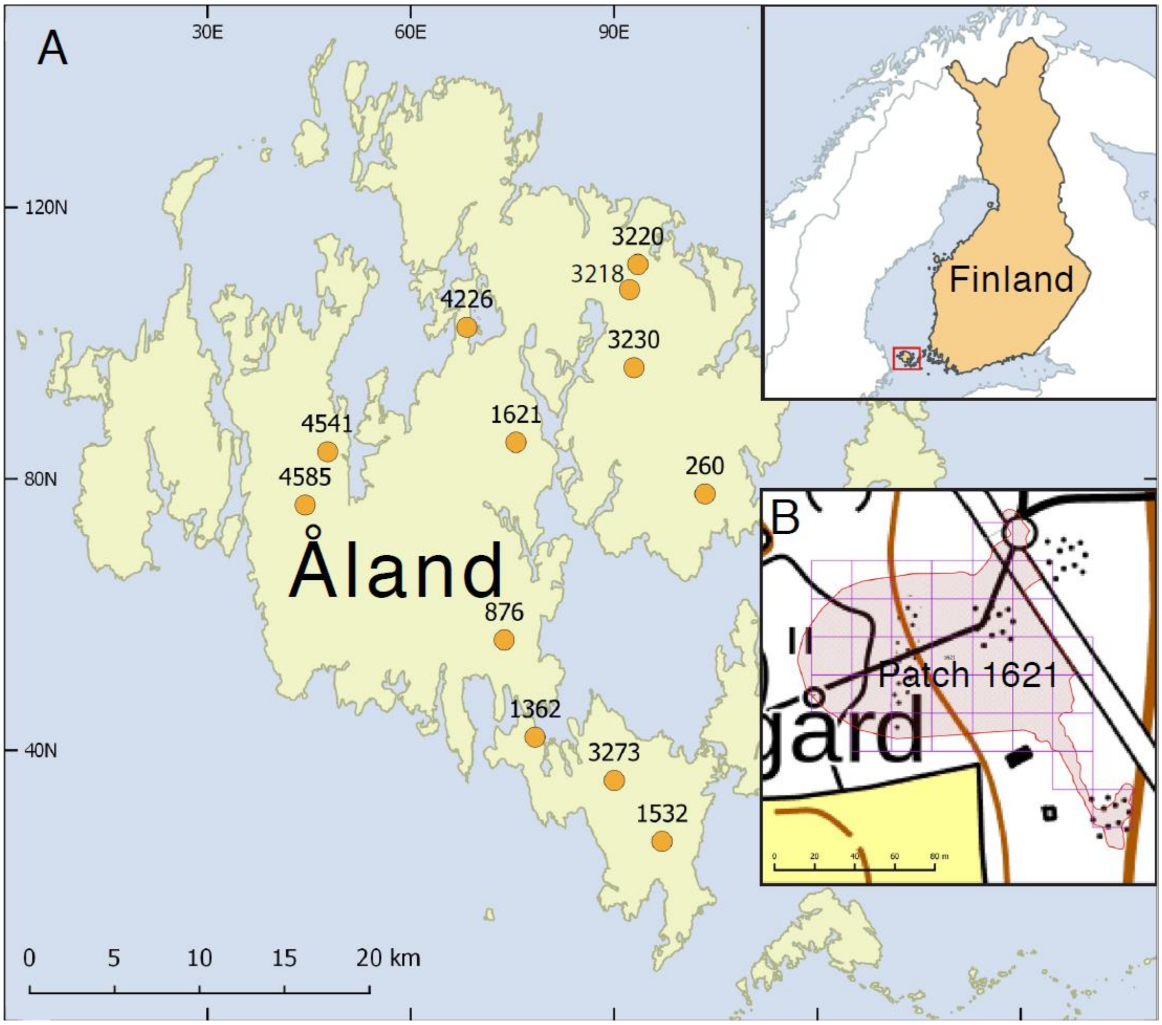

Appendix S2. Methods

*Field observations and data collection*

The initial patch selection was carried out based on the drought and abundance statistics of host plants recorded during the annual surveys (Ojanen et al. 2013), ensuring variability in drought stress yet stable host plant abundance to maintain several larval nests for observations (for more information about the population selection please see author's et al., in prep). As the field assessments were conducted every second week, we additionally disregarded habitat patches too close to human settlement.

For the drought stress we used a categorical measure; 1 = green and lush plants, 2 = still green but wilted plants, and 3 = yellow/brown plants with crispy and very dry leaves. We estimated the percentage of host plants within each category for measuring drought stress (*ds*) in each measurement along the summers, then we obtained one unique value per cell:

$$ds = [(1*0) + (2*50) + (3*100)]/ 100$$

, where one, two and three are the values estimated for each category. Values close to zero represented lush and green host plants, meanwhile values close to 1 represented dry host plants.

The terrain slope and aspect were calculated once using GDAL version 2.1 (2016) from the National Land Survey of Finland's 10 m elevation model (https://www.maanmittauslaitos.fi/en). The values were measured for each of the grid cells using their centroid coordinates. Both transformed field aspect (tasp) and the slope corrected transformed aspect (tasl) are bounded by -1 to 1, where -1 indicates a north-eastern facing slope and +1 indicates a south-western facing slope. The slope corrected transformed aspect is tasp weighted by the steepness of the slope.

To confirm the location of the nests within surveys, we compared locations of nests from summer and autumn surveys for 2015-2018. Except for 2018 where severe drought killed all but two nests by the end of the summer, there was little difference in the nest locations*. Then, we conducted microhabitat selection analysis using the summer data and autumn data for the same years and found no difference in results.

* Locations of nests in summer and autumn surveys for the years 2015-2018. Red indicates the presence of a nest within a particular cell (quadrat), with a darker shade indicating more nests. Grey indicates the absence of a nest in a particular cell. Results are divided by patch (right axis).

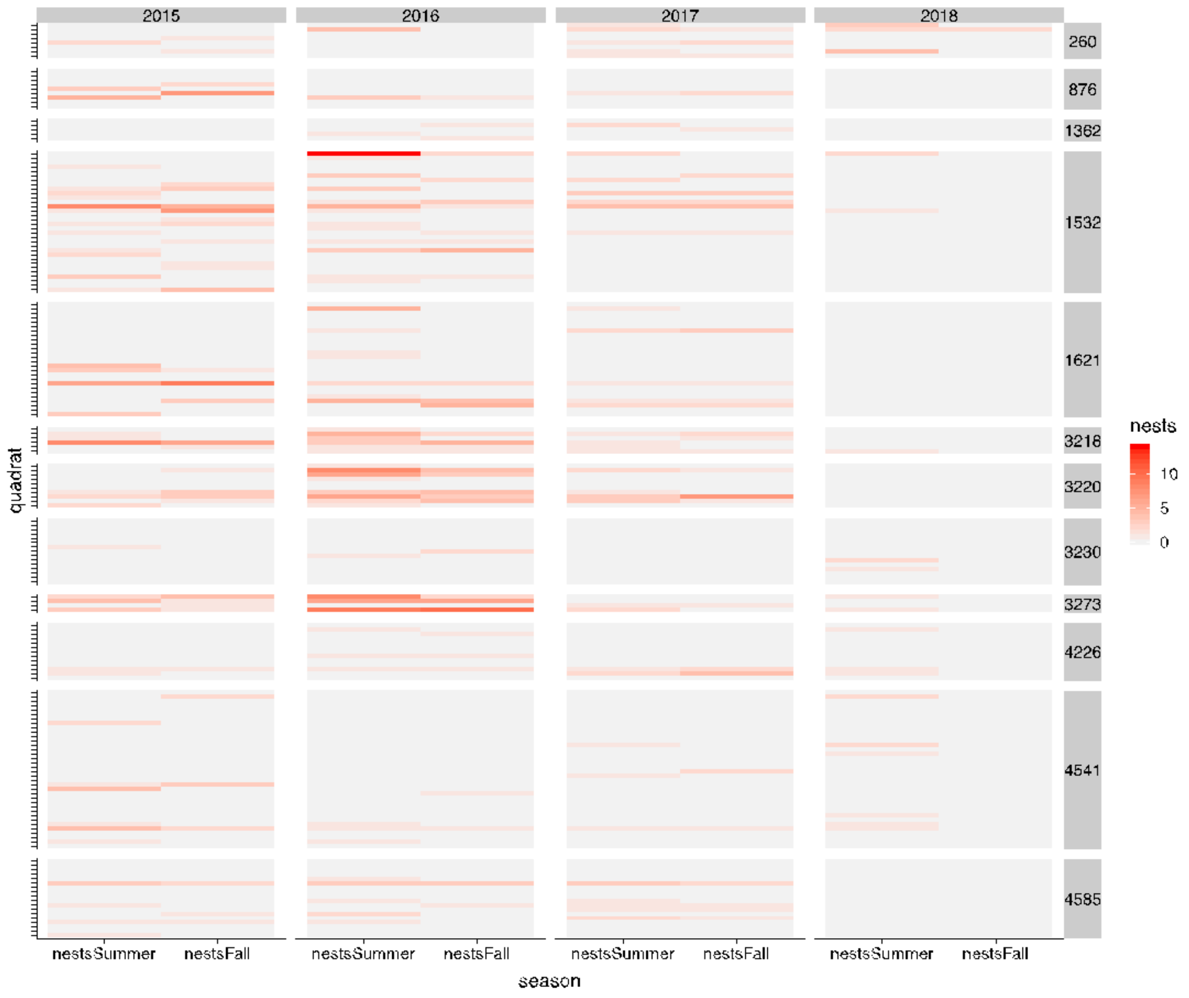

*Variation in host plant abundance and drought stress*

Beta regressions cannot accommodate zeros or ones in the response variable and so we transformed the proportion of host plant abundance and the proportion of drought stress using the following formula recommended by Smithson & Verkuilen*:

$$(y^*(n - 1) + 0.5)/n$$

, where $n$ is the number of samples. We implemented intercept-only beta models with cell nested within population nested within year as a random effect.

* Smithson M., & Verkuilen J. (2006). A better lemon squeezer? Maximum-likelihood regression with beta-distributed dependent variables. *Psychological Methods*, 11(1), 54-71.

Figure S3. Constructed mesh within the populations for speeding computing analysis. Red outlines are patch borders and open circles are centroids of 20x20 grid cells.

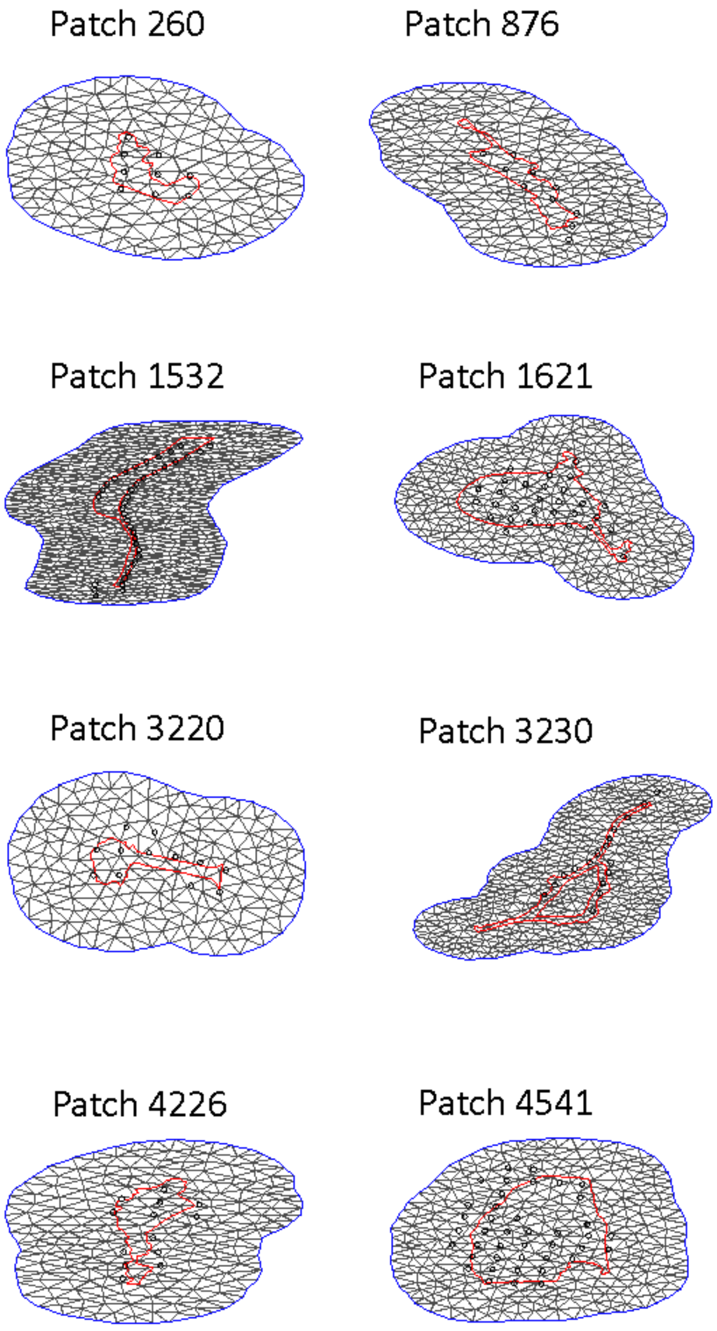

Table S4. Principal components (PC) retained after analysis.

| | PC1 | PC2 | PC3 |
|---|---|---|---|
| Eigenvalue | 1.65 | 1.22 | 1.11 |
| Proportion of variance | 0.28 | 0.20 | 0.18 |
| Cumulative variation | 0.28 | 0.48 | 0.66 |
| Variable loading | | | |
| Soil depth (cm) | 0.22 | 0.62 | -0.40 |
| Vegetation height (m) | 0.09 | 0.10 | 0.74 |
| Canopy coverage (%) | 0.37 | 0.59 | 0.02 |
| Terrain slope | 0.09 | -0.36 | -0.54 |
| Transformed field aspect (tasp) | -0.64 | 0.27 | -0.06 |
| Transformed weight of the slope (tasl) | -0.62 | 0.25 | -0.04 |

Figure S5. Boxplots showing variation in host plant abundance (A) and proportion of host plants showing drought stress (B), and the proportion of host plants showing drought stress during the period (C). Values in (A) and (B) were measured per cell four times per year except in 2018 where measurements were taken only in one time point, and values in (C) were taken from the time point during the driest summer period.

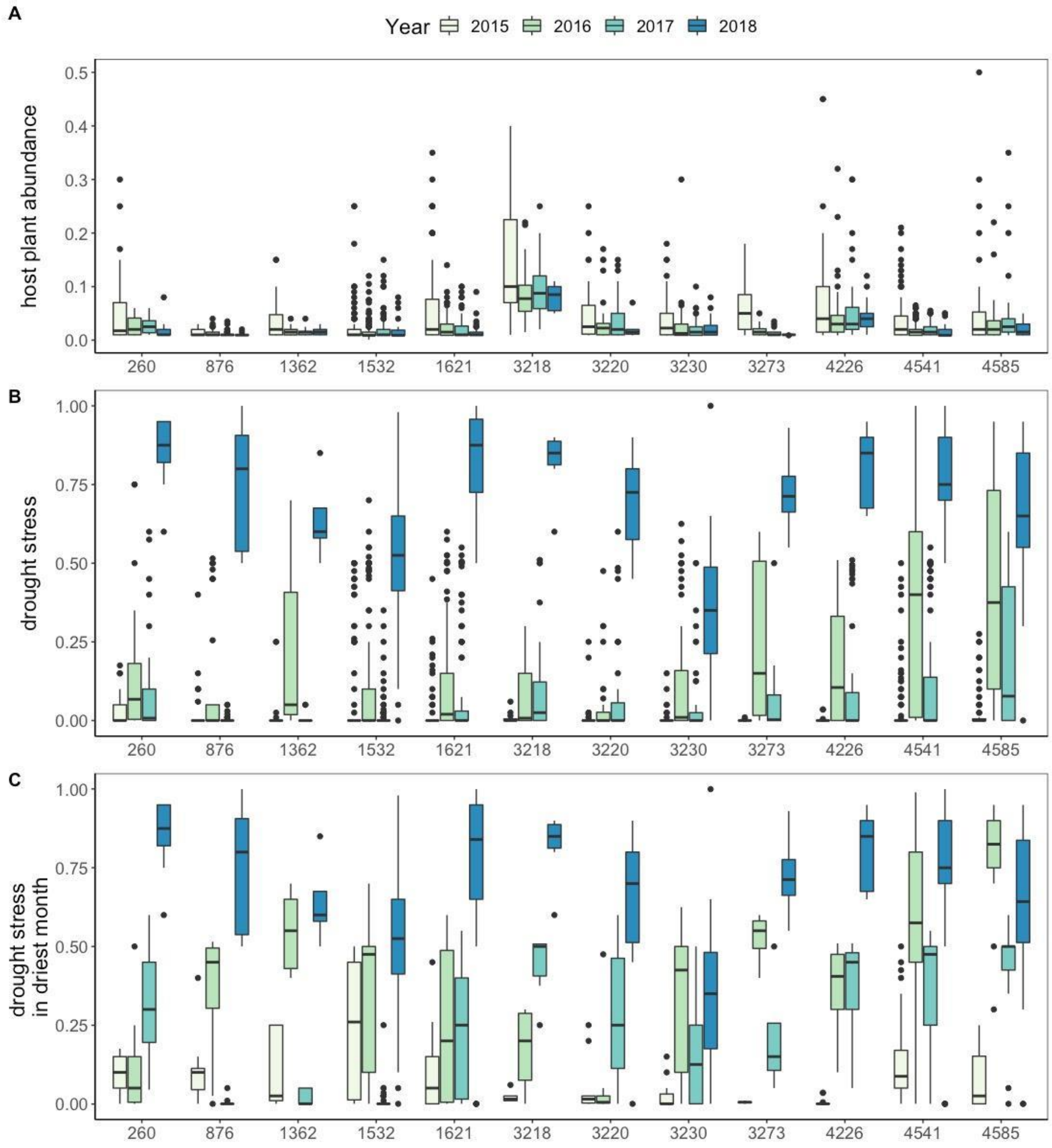

Figure S6. Histograms showing the yearly distributions of the proportion of host plants with signs of drought stress. Available but unoccupied microsites are shown in grey, and occupied sites are shown in blue.

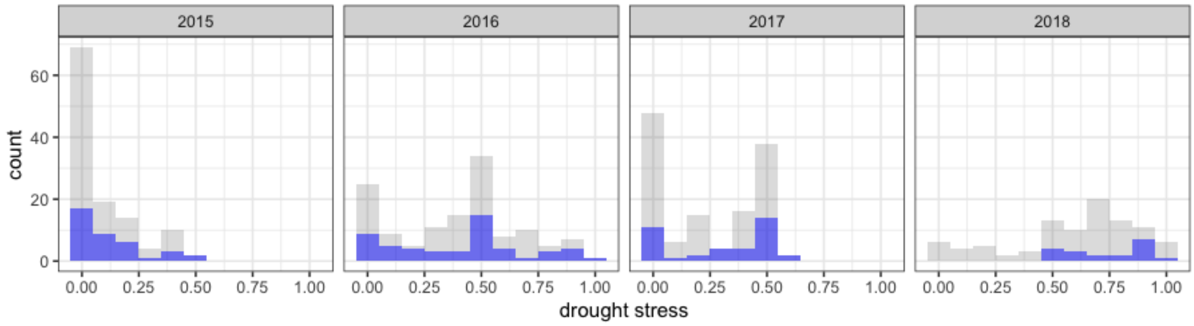

Figure S7. Heatmap showing pairwise Pearson correlations of host plant abundance measured at different time points (t1 = time point 1, t2 = time point 2, etc.) and years (2015-2018).

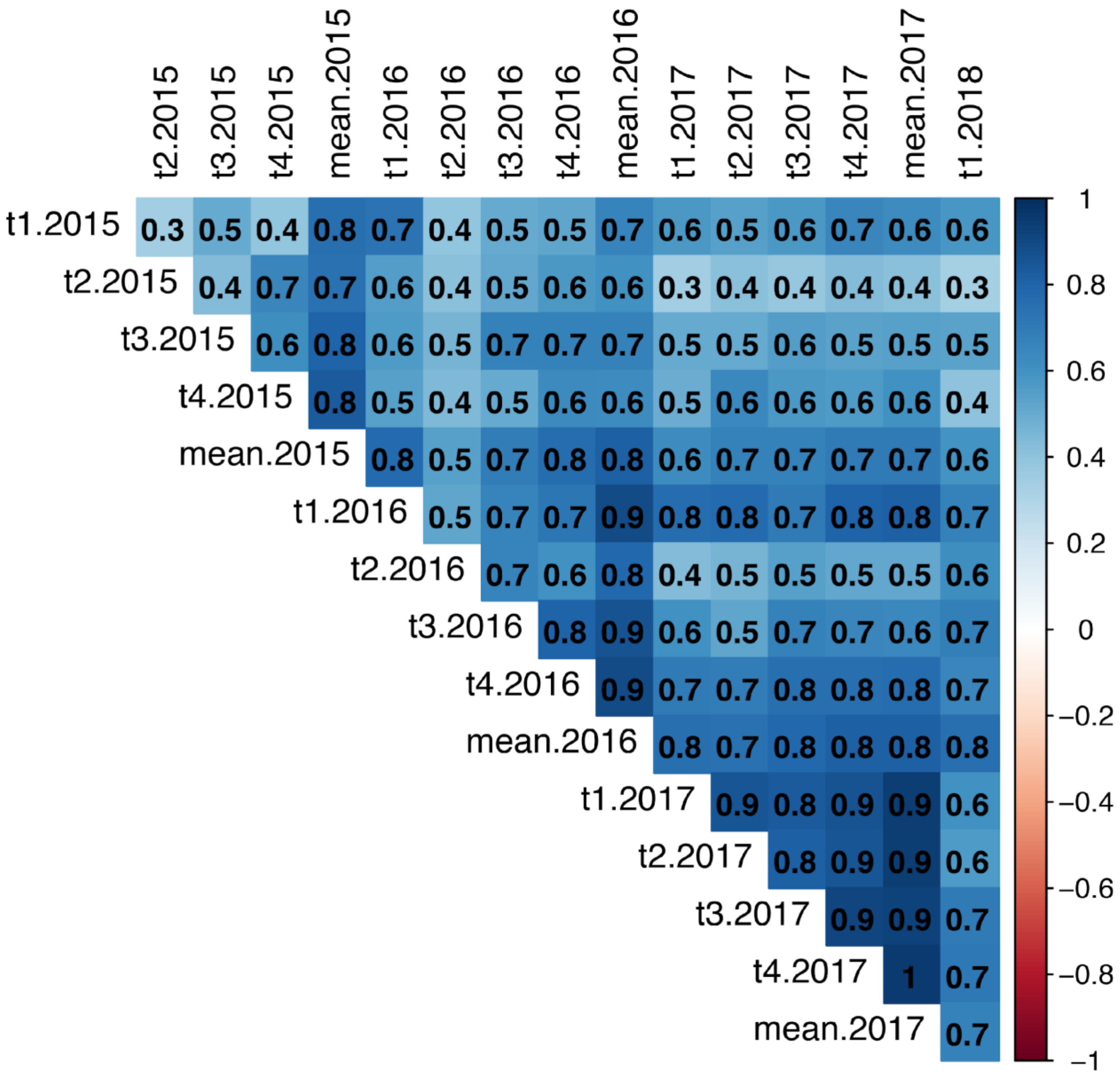

Figure S8. Heatmap showing pairwise Pearson correlations of proportion of host plants showing signs of drought stress measured at different time points (t1 = time point 1, t2 = time point 2, etc.) and years (2015-2018).

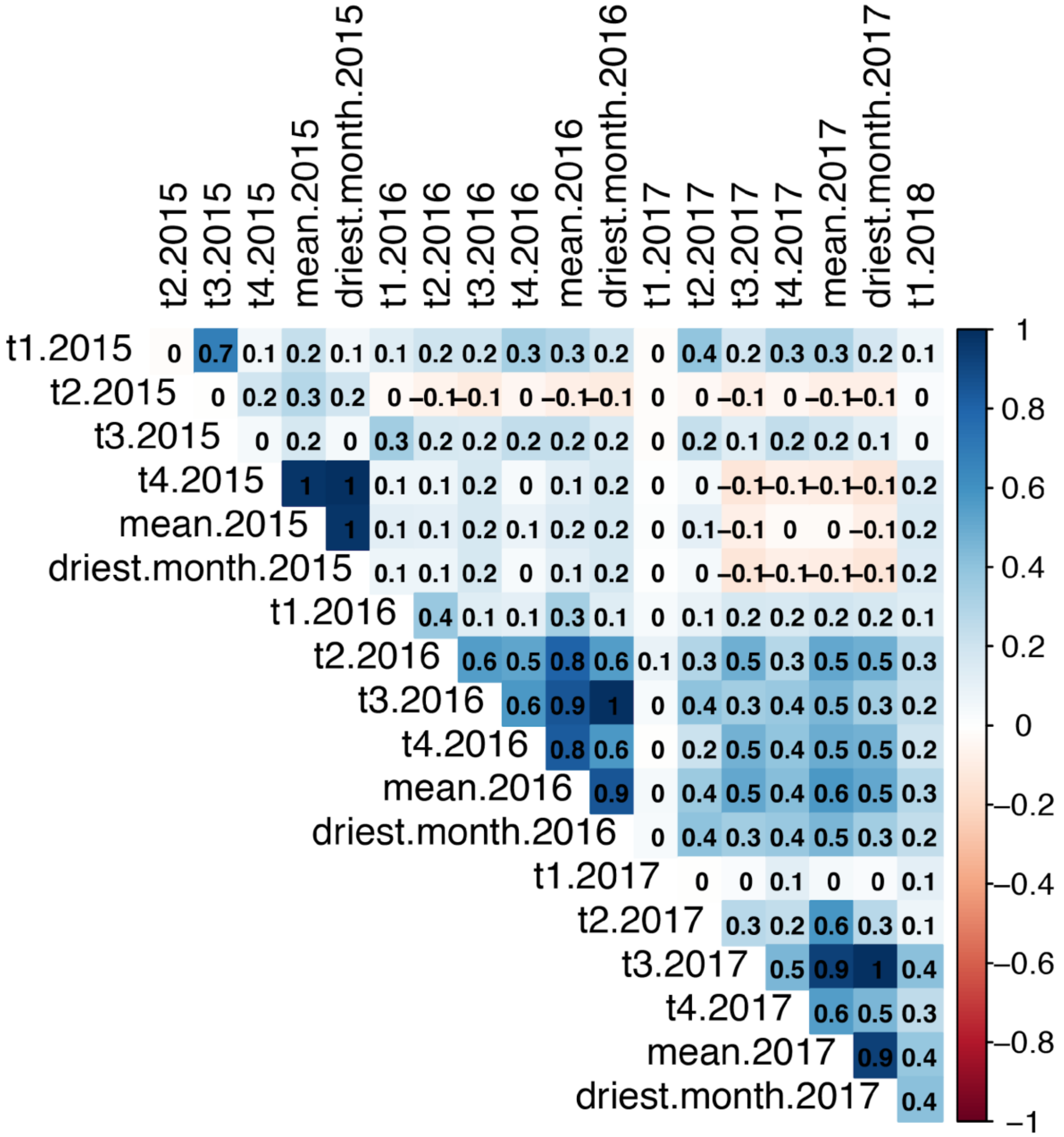

Figure S9. Pearson correlations among predictor variables included in the long-term binomial and hurdle models.

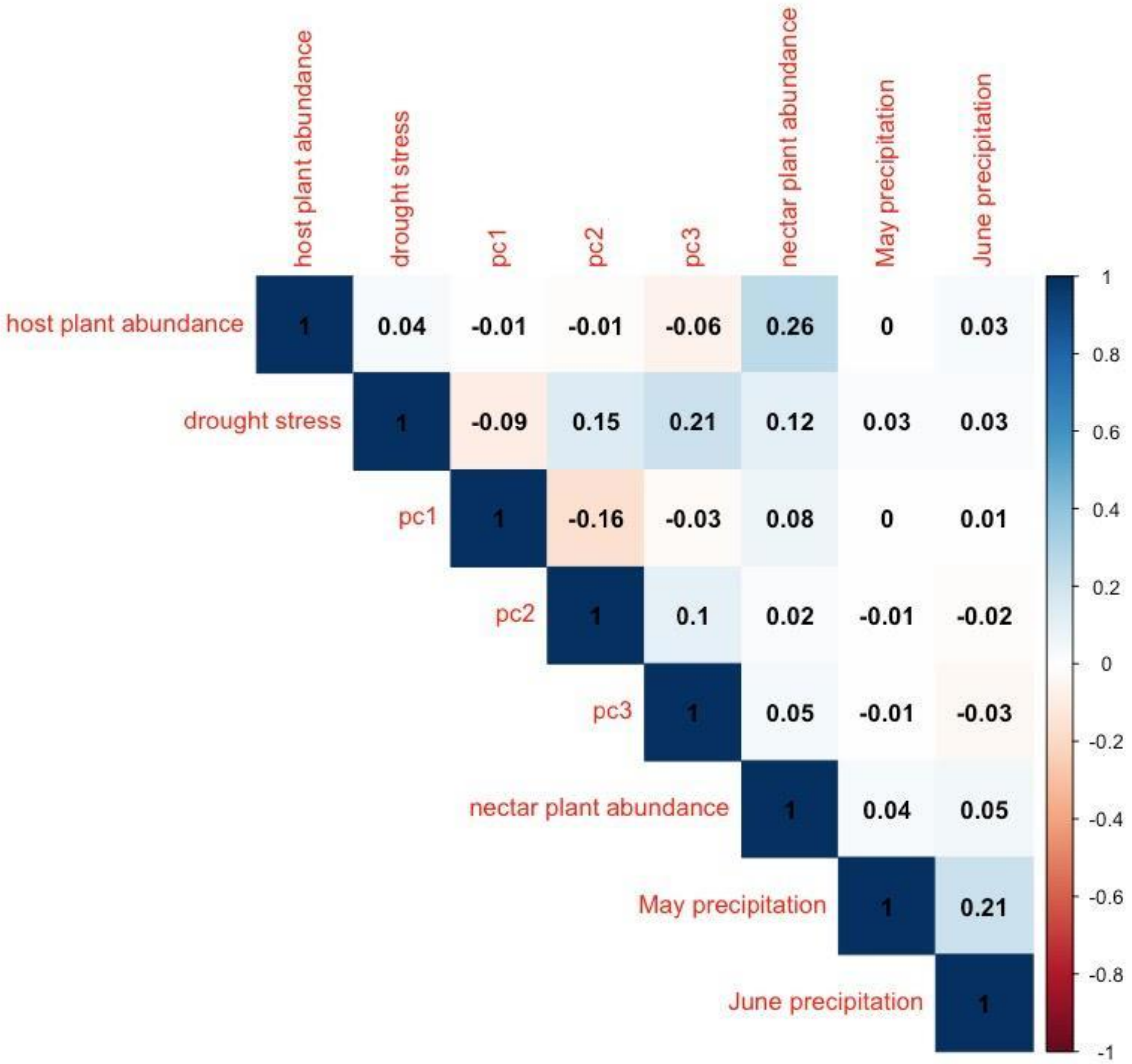

Table S10. Posterior mean estimates, standard deviations (SD) and 95% credibility intervals from hurdle INLA models using four-years of summer nest counts and ten-years of autumn nest counts. Host plant abundance was entered as a yearly mean value per cell in the four-year model, and as the mean across all years of field measurement per cell (2015-2018) for the ten-year model. Drought stress represents the proportion of host plants with signs of drought stress measured at the driest time point per cell per year for the four-year model, and the mean of these values across all years of field measurement was used for the ten-year model. Both models include a spatiotemporal random effect.

| | | four-year model | | | ten-year model | | |
|---|---|---|---|---|---|---|---|
| | Parameters | mean | SD | 95% credibility | mean | SD | 95% credibility |
| Zero-inflation probability | intercept | **-1.61** | **0.25** | **-2.15 , -1.15** | **-2.28** | **0.23** | **-2.76 , -1.86** |
| | host plant abundance | **0.78** | **0.13** | **0.53 , 1.05** | **0.73** | **0.12** | **0.51 , 0.98** |
| | drought stress | **0.39** | **0.15** | **0.11 , 0.68** | **0.71** | **0.15** | **0.42 , 1.01** |
| | abundance of nectar plants | 0.07 | 0.14 | -0.20 , 0.34 | 0.12 | 0.11 | -0.11 , 0.34 |
| | pc1 | **-0.42** | **0.13** | **-0.67 , -0.17** | **-0.40** | **0.11** | **-0.62 , -0.18** |
| | pc2 | **-0.47** | **0.15** | **-0.78 , -0.18** | **-0.52** | **0.14** | **-0.80 , -0.25** |
| | pc3 | **0.35** | **0.14** | **0.07 , 0.63** | 0.08 | 0.13 | -0.18 , 0.33 |
| | May precipitation | | | | -0.05 | 0.10 | -0.26 , 0.15 |
| | June precipitation | | | | **0.27** | **0.11** | **0.04 , 0.49** |
| | host plant abundance x May precip | | | | -0.07 | 0.08 | -0.23 , 0.09 |
| | drought stress x May precip | | | | **-0.21** | **0.10** | **-0.42 , -0.01** |
| | host plant abundance x June precip | | | | -0.03 | 0.09 | -0.20 , 0.15 |
| | drought stress x June precip | | | | 0.12 | 0.11 | -0.10 , 0.35 |
| Truncated Poisson | intercept | **-6.91** | **0.34** | **-7.65 , -6.32** | **-7.34** | **0.27** | **-7.91 , -6.83** |
| | host plant abundance | **0.23** | **0.08** | **0.07 , 0.39** | **0.28** | **0.10** | **0.09 , 0.48** |
| | drought stress | 0.07 | 0.14 | -0.21 , 0.35 | **0.62** | **0.16** | **0.33 , 0.94** |
| | abundance of nectar plants | -0.01 | 0.08 | -0.17 , 0.15 | -0.01 | 0.08 | -0.17 , 0.15 |
| | pc1 | **-0.31** | **0.13** | **-0.57 , -0.07** | -0.19 | 0.11 | -0.40 , 0.02 |
| | pc2 | -0.18 | 0.18 | -0.55 , 0.16 | **-0.36** | **0.14** | **-0.64 , -0.09** |
| | pc3 | 0.24 | 0.14 | -0.04 , 0.52 | **0.22** | **0.11** | **0.004 , 0.44** |
| | May precipitation | | | | -0.10 | 0.09 | -0.29 , 0.09 |
| | June precipitation | | | | **0.31** | **0.10** | **0.11 , 0.51** |
| | host plant abundance x May precip | | | | 0.02 | 0.05 | -0.08 , 0.13 |
| | drought stress x May precip | | | | 0.03 | 0.09 | -0.15 , 0.21 |
| | host plant abundance x June precip | | | | 0.13 | 0.08 | -0.02 , 0.28 |
| | drought stress x June precip | | | | 0.11 | 0.10 | -0.09 , 0.30 |
| | Random effects | | | | | | |
| | temporal correlation | 0.63 | 0.13 | 0.34 , 0.83 | 0.76 | 0.05 | 0.65 , 0.85 |
| | variance of spatial random effect | 1.06 | | | 1.63 | | |

Table S11. Posterior mean estimates, standard deviations (sd) and 95% credibility intervals from spatiotemporal binomial INLA models on nest presence using a mix of autumn and summer nest locations (autumn for 2015-17, summer for 2018; n = 602), summer nest locations (2015-2018; n = 617), and autumn nest locations (2015-17; n = 490).

| | mix | | | summer nests | | | autumn nests | | |
|---|---|---|---|---|---|---|---|---|---|
| Parameter | mean | sd | 95% credibility | mean | sd | 95% credibility | mean | sd | 95% credibility |
| intercept | **-1.8** | **0.26** | **-2.4 , -1.3** | **-1.5** | **0.23** | **-2.0 , -1.04** | **-1.8** | **0.29** | **-2.4 , -1.2** |
| host plant abundance | **0.79** | **0.13** | **0.54 , 1.07** | **0.78** | **0.13** | **0.53 , 1.04** | **0.81** | **0.15** | **0.53 , 1.11** |
| drought stress | **0.39** | **0.15** | **0.10 , 0.69** | **0.34** | **0.15** | **0.06 , 0.65** | **0.43** | **0.15** | **0.14 , 0.74** |
| nectar plant abundance | 0.18 | 0.14 | -0.09 , 0.47 | 0.10 | 0.13 | -0.15 , 0.36 | 0.22 | 0.15 | -0.05 , 0.52 |
| pc1 | **-0.38** | **0.14** | **-0.66 , -0.12** | **-0.42** | **0.12** | **-0.65 , -0.19** | **-0.31** | **0.14** | **-0.60 , -0.04** |
| pc2 | **-0.51** | **0.17** | **-0.85 , -0.20** | **-0.47** | **0.14** | **-0.75 , -0.21** | **-0.43** | **0.17** | **-0.77 , -0.09** |
| pc3 | 0.19 | 0.15 | -0.11 , 0.49 | 0.28 | 0.13 | 0.02 , 0.54 | 0.18 | 0.16 | -0.14 , 0.49 |
| Random effects | | | | | | | | | |
| temporal correlation | 0.69 | 0.23 | 0.11 , 0.96 | 0.44 | 0.22 | -0.04 , 0.80 | 0.80 | 0.25 | 0.06 , 0.99 |
| variance of spatial random effect | 0.16 | | | 0.13 | | | 0.12 | | |

Figure S12. Relationships between precipitation and mean temperature in May (A) and June (B). Points represent populations, and lines show linear trends within years.

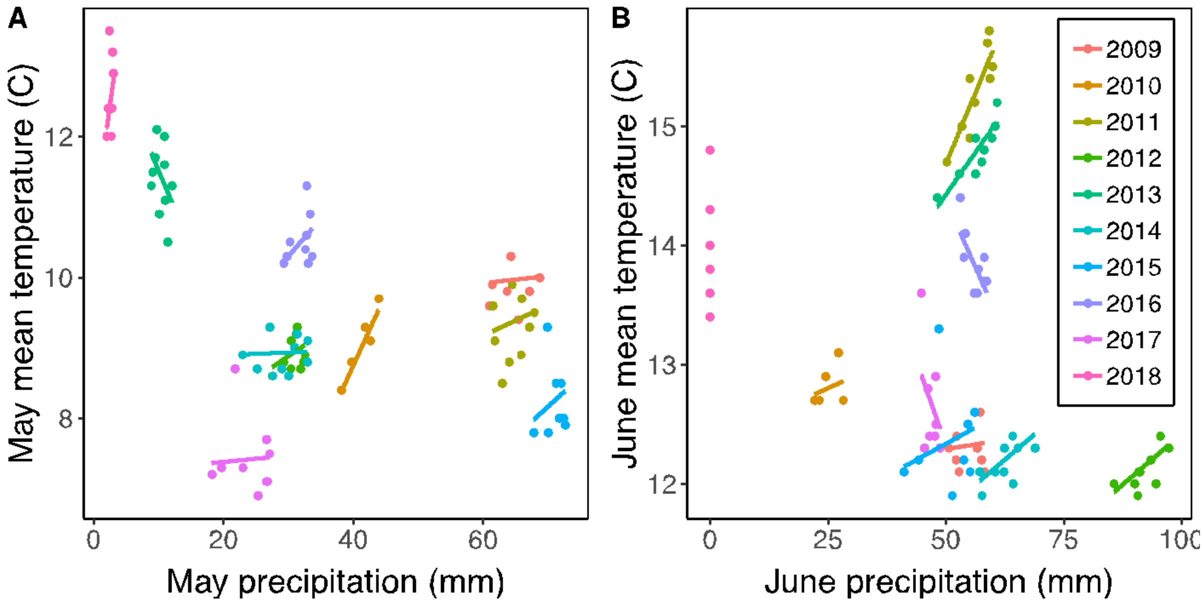

Figure S13. Histograms showing mean drought stress values of available but unused microhabitats (grey), used microhabitats where nests survived overwinter (green), and used microhabitats where nests died overwinter (red). Used microhabitats were more likely to have higher mean drought stress values, and nests were more likely to survive in these microhabitats.

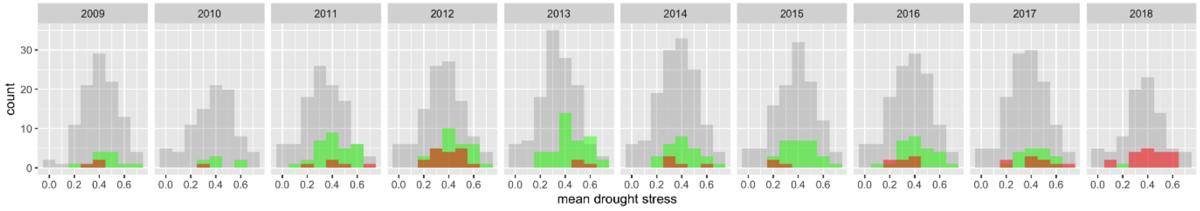
